# CRISPR-bind: a simple, custom CRISPR/dCas9-mediated labeling of genomic DNA for mapping in nanochannel arrays

**DOI:** 10.1101/371518

**Authors:** Denghong Zhang, Saki Chan, Kenneth Sugerman, Joyce Lee, Ernest T. Lam, Sven Bocklandt, Han Cao, Alex R. Hastie

**Affiliations:** Bionano Genomics, San Diego, CA

**Keywords:** dCas9, tracrRNA, CRISPR, Custom labeling, Targeted labeling, Optical mapping, Genome mapping

## Abstract

Bionano genome mapping is a robust optical mapping technology used for *de novo* construction of whole genomes using ultra-long DNA molecules, able to efficiently interrogate genomic structural variation. It is also used for functional analysis such as epigenetic analysis and DNA replication mapping and kinetics. Genomic labeling for genome mapping is currently specified by a single strand nicking restriction enzyme followed by fluorophore incorporation by nick-translation (NLRS), or by a direct label and stain (DLS) chemistry which conjugates a fluorophore directly to an enzyme-defined recognition site. Although these methods are efficient and produce high quality whole genome mapping data, they are limited by the number of available enzymes—and thus the number of recognition sequences—to choose from. The ability to label other sequences can provide higher definition in the data and may be used for countless additional applications. Previously, custom labeling was accomplished via the nick-translation approach using CRISPR-Cas9, leveraging Cas9 mutant D10A which has one of its cleavage sites deactivated, thus effectively converting the CRISPR-Cas9 complex into a nickase with customizable target sequences. Here we have improved upon this approach by using dCas9, a nuclease-deficient double knockout Cas9 with no cutting activity, to directly label DNA with a fluorescent CRISPR-dCas9 complex (CRISPR-bind). Unlike labeling with CRISPR-Cas9 D10A nickase, in which nicking, labeling, and repair by ligation, all occur as separate steps, the new assay has the advantage of labeling DNA in one step, since the CRISPR-dCas9 complex itself is fluorescent and remains bound during imaging. CRISPR-bind can be added directly to a sample that has already been labeled using DLS or NLRS, thus overlaying additional information onto the same molecules. Using the dCas9 protein assembled with custom target crRNA and fluorescently labeled tracrRNA, we demonstrate rapid labeling of repetitive DUF1220 elements. We also combine NLRS-based whole genome mapping with CRISPR-bind labeling targeting Alu loci. This rapid, convenient, non-damaging, and cost-effective technology is a valuable tool for custom labeling of any CRISPR-Cas9 amenable target sequence.

## Introduction

Bionano Genomics’ optical mapping technology generates *de novo* assembled genome maps which can be used to assemble genome sequences, to analyze structural variation, and to study various biological processes. Native ultra-long DNA is isolated from cells or organisms, a specific sequence motif is labeled, the labeled DNA is linearized and imaged in nanochannel arrays, and *de novo* maps of the entire genome generated by pairwise alignment of the imaged molecules. These maps can scaffold sequence contigs and generate structurally accurate genome assemblies for human, plant and animal genomes, including the 32 Gbp axolotl genome (Nowoshilow 2018). Bionano maps enable the analysis of human structural variation and is highly accurate and sensitive for detection of SVs >500 bp which are typically missed by sequencing based methods, including balanced events that are invisible to standard cytogenetic techniques (Hastie 2017, Barseghyan 2017, Chaisson 2018). Finally, whole genome mapping is being used for functional analysis of various biological processes such as DNA replication (Klein 2017) and epigenetic modifications (Grunwald 2017, Tslil 2018).

Current labeling chemistries for Bionano genome mapping involve either nick translation using nick-label-repair-stain chemistry (NLRS) or direct labeling using the direct label and stain chemistry (DLS) (both from Bionano Genomics). These chemistries are extremely robust, but a limited number of enzymes (and thus target sequences) which can be used in NLRS and DLS chemistries are available. The ability to easily generate custom targeted tools would open the platform to countless new applications.

The type II clustered regularly interspaced short palindromic repeats (CRISPR)-associated Cas9 system derived from *Streptococcus pyogenes* has become the dominant tool for targeted genome editing (Sternberg 2015) because of its target sequence customizability and high binding and enzymatic efficiency. To achieve site-specific DNA recognition and cleavage, the protein, Cas9, forms a ribonucleoprotein (RNP) complex with guide RNA (gRNA). gRNA is a duplex consisting of CRISPR RNA (crRNA) and trans-activating crRNA (tracrRNA). The specificity comes from the crRNA, which contains an approximately 20-nucleotide sequence, complementary to the target sequence; the remainder of the crRNA is partially complementary to the tracrRNA, facilitating annealing. Once the RNP binds to the target sequence, the HNH and RuvC-like nuclease domains of Cas9 generate double-stranded breaks (Jinek 2012, Gasiunas 2012).

CRISPR-Cas9 is a double-strand DNA endonuclease by default; however, it has been modified to function as a nickase, methyltransferase, transcription activator or repressor, and labeling tool (Lo 2017). Cas9 nuclease-deficient derivatives (dCas9) are also used for control of gene expression (Qi 2013, Dominguez 2016) and visualization of genomic loci in cells and tissues (Deng 2015) through fusion with a fluorescent protein (Chen 2013).

McCaffrey et al. (McCaffrey 2016) has applied single-strand nicking CRISPR-Cas9 D10A to specify a target for nick-translation of fluorescent nucleotides, in order to label DNA for analysis in nanochannel arrays on Bionano Saphyr. They have been able to mark tandem repeat arrays of DUF1220, translocation breakpoints, and other targets. They have also combined targeted CRISPR-Cas9 labeling with Nt.BspQI nick-labeling to map the CRISPR-Cas9 D10A encoded targets to the human reference, including characterization of telomere sequences (McCaffrey 2017).

Here we have leveraged the CRISPR-dCas9 system for custom labeling of bacterial artificial chromosome (BAC) DNA and human genomic DNA for analysis in nanochannel arrays on the Bionano Genomics Saphyr system. The labeling method is analogous to the method developed by McCaffrey et al. but instead of the multistep process of nick, label, repair (ligation), it is a simple, single-step reaction in which fluorescent RNP is bound to the DNA target sites. We show that this method, CRISPR-bind labeling, is able to efficiently label unlabeled and previously labeled BAC and human genomic DNA.

## Results

In order to develop a simple, fast and cost-effective method to custom label DNA with a fluorescent probe that can be measured in nanochannel arrays, we have developed the CRISPR-bind method. In this method, a guide RNA (gRNA) is made by annealing a fluorescently conjugated tracrRNA to a crRNA that contains a ~20 nucleotide sequence complementary to the target site. RNP complexes are then formed by combining dCas9 protein and the prepared gRNA (Figure 1). The double mutations in the HNH and RuvC-like nuclease domain of dCas9 remove catalytic activity while maintaining very high specificity and efficient binding (Qi 2013).

**Figure 1.**
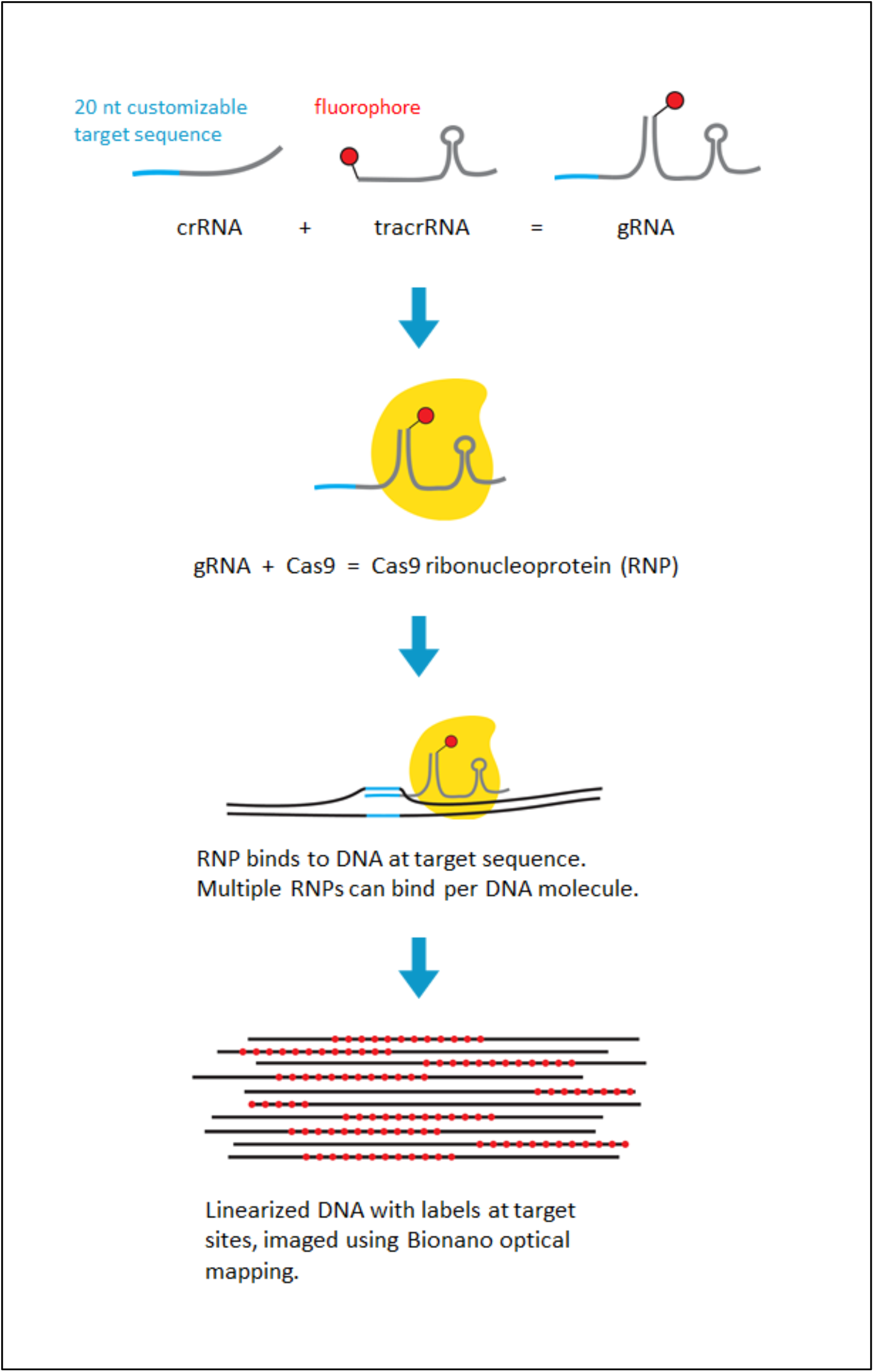
Schematic of CRISPR-bind genome mapping.

With dCas9, fluorescently labeled tracrRNA, and a custom crRNA probe, we can target any sequence amenable to CRISPR-Cas9 targeting by modifying only the crRNA probe. We have applied this DNA labeling method, CRISPR-bind, to labeling BAC DNA and megabase-length human genomic DNA, successfully introduced the DNA into nanochannel arrays, and imaged them using Bionano Irys or Saphyr systems (Figure 1).

### Labeling DUF1220 repeats on BAC DNA by CRISPR-bind

We first targeted CRISPR-bind to the DUF1220 repeat present on chromosome 1. In the human genome, this sequence is present in variable copy number and organized in several tandem arrays of the repeat unit; the copy number has been correlated with brain size in primates (Dumas 2012). The exact lengths of long tandem repeat arrays are extremely challenging to measure since they are longer than sequence reads and too short to visualize by FISH or other cytogenetic methods. Optical mapping of ultra-long molecules that span the repeat arrays allows the repeat units to be counted and the arrays placed into context; however, the repeats are typically not marked by current labeling enzymes.

We synthesized a crRNA targeting DUF1220 monomers (based on McCaffrey et al., 2016) and assembled a CRISPR-bind RNP complex with fluorescently labeled tracrRNA. The complex was incubated with linearized BAC DNA, anticipated to contain 11 copies of DUF1220. CRISPR-bind complex concentration was titrated to find the optimal concentration of 15 nM (data not shown). At this concentration, the signal-to-noise ratio (SNR) allowed the sample to be loaded directly onto the Saphyr chip without having to clean up excess fluorophores. From the images we measured the efficiency of detection of labels at each binding site as well as the frequency that a signal was called outside of the expected DUF1220 loci (Figure 2). Target labeling efficiency was 87% while 16% of labels appeared to be off-target. This performance is similar to standard nick-labeling with Nt.BspQI.

**Figure 2.**
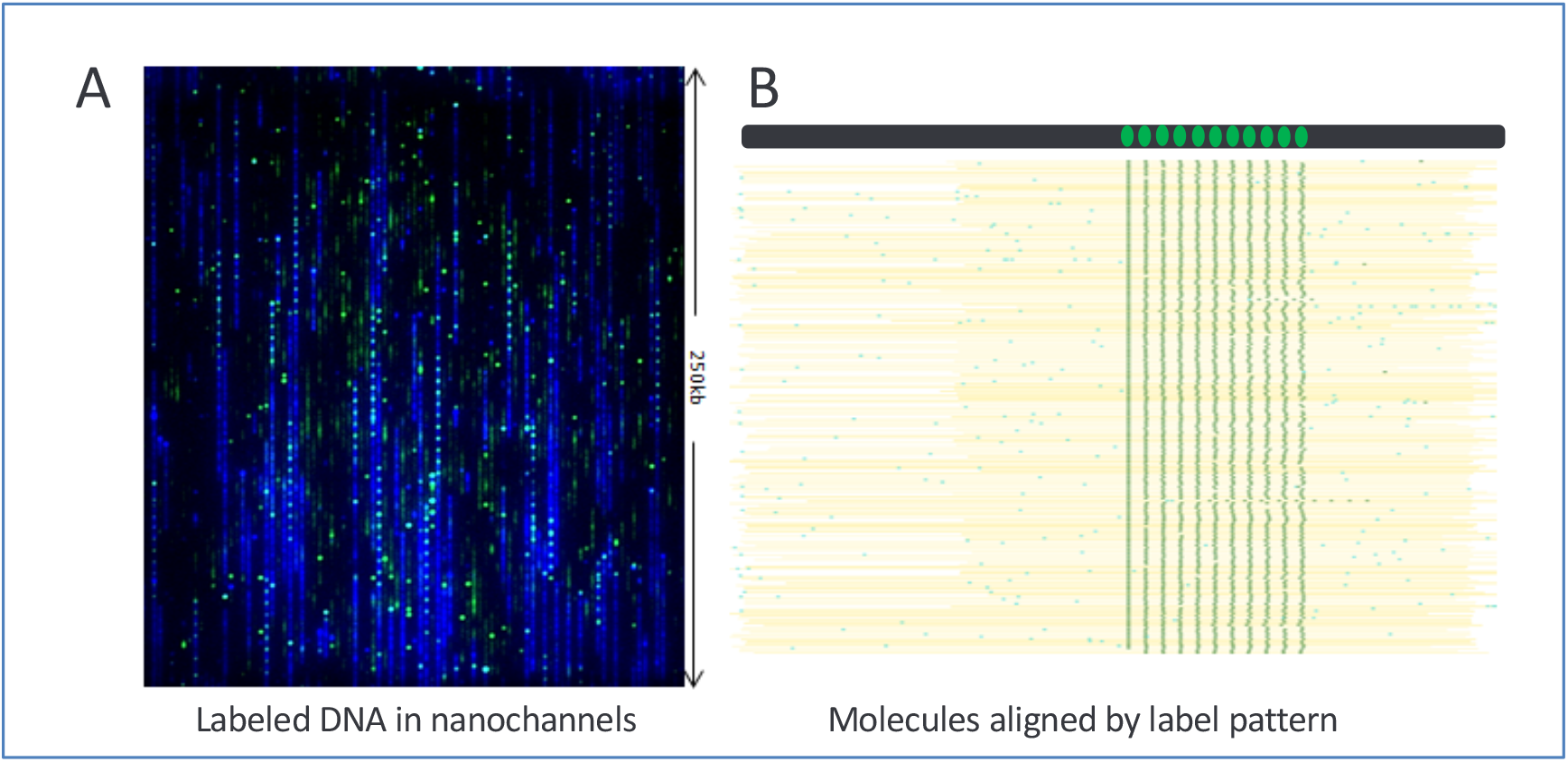
CRISPR-bind (DUF1220) labeling on a BAC Clone. (**A**) The CRISPR-bind labeling method was used to label each DUF1220 repeat unit using a gRNA designed to be complimentary to the DUF1220 repeat unit. The resulting tandem repeat labels (shown in green) on the BAC DNA (its backbone shown in blue) were separated by approximately 4.7 kbp as predicted. (**B**) The single molecule pile-up of molecules containing the DUF1220 specific labels is shown.

### CRISPR-bind labeling combined with nick-labeling to define and localize Alu sequences

In order to test CRISPR-bind for specificity and efficiency on human genomic DNA, we chose to use a target sequence which binds to about a quarter of all Alu repeats across the genome. Alu repeats are the most abundant transposable elements in the human genome, and they can copy and insert themselves into new locations, sometimes altering gene expression and even causing disease (Deininger 1999). Unlike other potential CRISPR ^~^20-bp targets, this Alu repeat occurs very frequently in the genome, allowing for relatively high density of labels across the genome. Utilization of CRISPR-bind-Alu can add 70% additional labels to a standard genome mapping experiment compared to nick-labeling alone, or about 50% more compared to DLS using DLE-1 alone, potentially providing higher information density for genomic variation analysis. Also, since the CRISPR-bind can be an alternate color, the second color signal can provide additional pattern uniqueness.

In order to estimate specificity and efficiency of binding and detection, we first labeled Nt.BspQI sites with standard NLRS labeling using a green dye, followed by labeling with Alu-targeting CRISPR-bind complex using a red dye on a human pseudo-haploid hydatidiform mole cell line, CHM1. Data was collected and mapped based on the Nt.BspQI pattern to the human reference (Figure 3). Nt.BspQI sites aligned to the *in silico* digestion of hg19, and the Alu/CRISPR-bind sites aligned with each other, indicating that the binding was both specific and efficient. Alu specifying signal often corresponded to annotated Alu target sites but frequently a consensus of molecules showed labels inconsistent to the reference loci, which is consistent with the high degree of variation of Alu patterns from individual to individual (Batzer 1996).

**Figure 3.**
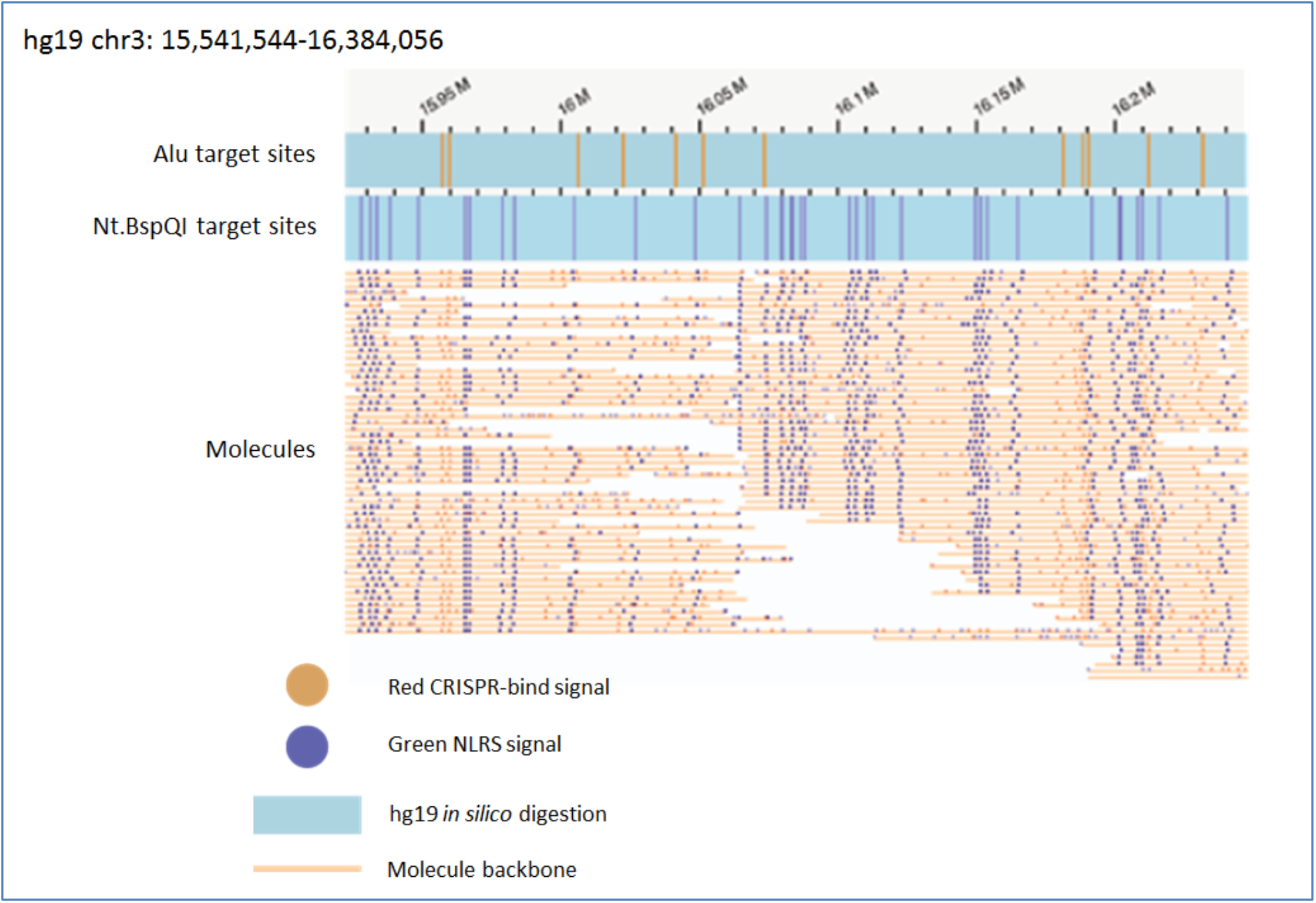
Alu repeat mapping on CHM1 by dCas9/Alu. Molecules labeled with both NLRS and CRISPR-bind, aligned by their nick-label patterns to an *in silico* digestion of hg19 reference with Nt.BspQI. Predicted Alu target sites are shown above. Nt.BspQI signal is shown on the molecules in purple and Alu signal is shown in orange. Approximately 90% of all molecules mapping to a particular region contain the same orange signal near a mapped nick-label. The Alu repeat signal forms a pattern concordant with predicted Alu sites

## Discussion

We have developed a new sequence-specific labeling method with dCas9 protein for use in optical mapping. This single-step labeling does not damage the DNA, and the flexible and efficient fluorescent tagging of specific sequences enables acquisition of context-specific sequence information, when performing single-molecule optical mapping in the Bionano Genomics nanochannel array. Not only can this new method improve the quality and sensitivity of whole-genome structural variation analysis by adding a second color and increasing information density, it is also able to target a wide variety of sequences such as long tandem repeats, viral integration sites, transgenes, and could even be used to genotype single nucleotide variants. Thus, this method can improve both automated high-throughput genome-wide mapping as well as targeted analyses of complex regions containing repetitive and structurally variant DNA.

## Materials and Methods

The CRISPR-Cas9 labeling methods presented here are largely based on McCaffrey et al. 2016.

### DNA purification and nick-labeling

Ultra-high molecular weight DNA was prepared from BAC clones CH17-353B19 (Invitrogen), and human cell line CHM1 (ref). Briefly, using the Bionano plug lysis protocol and reagents, cells were immobilized in low melting point agarose, lysed, and treated with Proteinase K, before being washed, solubilized with Agarase (Thermo Fisher), and drop-dialyzed.

The human genomic DNA sample (CHM1) was nick-labeled using the Bionano NLRS kit reagents and protocols. Briefly, DNA was digested with nickase Nt.BspQI (NEB), labeled with Taq DNA Polymerase (NEB) using fluorescent green nucleotides, and repaired with Taq DNA Ligase (NEB). The DNA backbone staining step was delayed until after CRISPR-binding.

### RNA design and preparation

The 20 nt crRNA target sequence was designed using the web server at crispr.mit.edu. DUF1220 crRNA (AAGUUCCUUUUAUGCAUUGG), and Alu crRNA (UGUAAUCCCAGCACUUUGGG) were synthesized by Synthego. Once received, the stock was diluted in TE Buffer to a working concentration of 100 μM.

Universal ATTO 647 labeled tracrRNA (Alt-R CRISPR-Cas9 tracrRNA) was purchased from Integrated DNA Technologies. The stock was diluted in TE Buffer to a working concentration of 100 μM.

### Guide RNA (gRNA) preparation

The fluorescent tracrRNA was annealed to Alu and DUF1220 crRNAs (separate reactions) by combining 500 pmol crRNA and 500 pmol tracrRNA in 1X NEBuffer3 and 1X BSA (NEB), in a total volume of 20 μL, and incubating at 4°C for 30 minutes.

### Fluorescent CRISPR-dCas9 ribonucleoprotein (RNP) assembly

Double knockout dCas9 protein D10A/H840A (no cleavage activity) was purchased from PNA Bio (cat #CD01) and was diluted to a stock concentration of 1 mg/mL or 6 μM with the provided diluent.

2 μL of the annealed gRNA (50 pmol) was mixed with 600 ng of dCas9 (3.6 pmol), in 1X NEBuffer 3 and 1X BSA (NEB) in a final volume of 10 μL, and incubated for 60 min at 37°C. The resulting RNP was expected to be 360 nM assuming 100% binding efficiency of dCas9 to gRNA (since gRNA was in excess). The RNP was stored at 4°C and used within 4 weeks.

### RNP binding of DNA and DNA backbone staining

150 ng of prepared DNA (from BACs and human, separately) was combined with 0.4 μL of RNP (15 nM final concentration of RNP) and incubated at 37°C for 60 min. The DNA backbone was stained using DNA Stain from the Bionano NLRS kit. The sample was stored at room temperature overnight and moved to 4°C for long-term storage.

### Data acquisition and visualization

DUF1220 mapping: 16 μL of the sample was loaded into an Irys Chip flow cell. An Irys run was initiated and after 20 cycles of electrophoresis and imaging, molecules were aligned to the reference sequence using IrysView.

Alu repeat mapping: 19 μL of the sample was loaded into a Saphyr Chip flow cell. A Saphyr run was initiated through Bionano Access software. After 30 cycles of electrophoresis and imaging, a *de novo* assembly was performed by Bionano Solve software. Consensus maps and molecules were aligned to the reference sequence and visualized using Bionano Access.

## References

Barseghyan H, Tang W et al. (2017) Next-generation mapping: a novel approach for detection of pathogenic structural variants with a potential utility in clinical diagnosis. Gen Med 2017. 9:90

Batzer MA, Arcot SS et al. (1996) Genetic variation of recent Alu insertions in human populations. J Mol Evol 42: 22

Chaisson MJP, Sanders AD et al. (2018) Multi-platform discovery of haplotype-resolved structural variation in human genomes. bioRxiv 193144; doi: https://doi.org/10.1101/193144

Chen B, Gilbert LA, Cimini BA, et al. (2013) Dynamic Imaging of Genomic Loci in Living Human Cells by an Optimized CRISPR/Cas System. Cell. 2013;155(7):1479–1491

Deininger PL, Batzer MA. (1999) Alu repeats and human disease. Mol Genet Metab. 1999 Jul;67(3):183–93. Review.

Deng, W, Shi X, Tjian R, Lionnet T, Singer. (2015) CASFISH: CRISPR/Cas9-mediated in situ labeling of genomic loci in fixed cells. Proc. Natl. Acad. Sci., 112 (38), 11870–11875

Dominguez AA, Lim WA, Qi LS.(2016) Beyond editing: repurposing CRISPR-Cas9 for precision genome regulation and interrogation. Nat Rev Mol Cell Biol. 2016 Jan; 17(1):5–15

Dumas, LJ et al. (2012) DUF1220-Domain Copy Number Implicated in Human Brain-Size Pathology and Evolution. American Journal of Human Genetics 91.3: 444–454

Gasiunas G, Barrangou R, Horvath P, Siksnys V. (2012) Cas9-crRNA ribonucleoprotein complex mediates specific DNA cleavage for adaptive immunity in bacteria. Proc. Natl. Acad. Sci. 109, E2579–E2586

Grunwald A, Sharim H et al. (2017) Reduced representation optical methylation mapping (R2OM2). bioRxiv 108084; doi: https://doi.org/10.1101/108084

Hastie AR, Lam ET et al. (2017) Rapid Automated Large Structural Variation Detection in a Diploid Genome by NanoChannel Based Next-Generation Mapping. bioRxiv 102764; doi: https://doi.org/10.1101/102764

Jinek M, Chylinski K, Fonfara I, Hauer M, Doudna JA, Charpentier E. (2012) A programmable dual-RNA-guided DNA endonuclease in adaptive bacterial immunity. Science 337, 816–821

Klein K, Wang W et al. (2017) Genome-Wide Identification of Early-Firing Human Replication Origins by Optical Replication Mapping. bioRxiv 214841; doi: https://doi.org/10.1101/214841

Lo A and Qia L. (2017) Genetic and epigenetic control of gene expression by CRISPR–Cas systems. Version 1. F1000Res. 2017; 6: F1000 Faculty Rev-747. doi: 10.12688/f1000research.11113.1

McCaffrey J, Sibert J, Zhang B, Zhang Y, Hu W, Riethman H, and Xiao M. (2016) CRISPR-CAS9 D10A nickase target-specific fluorescent labeling of double strand DNA for whole genome mapping and structural variation analysis. Nucleic Acids Res. Jan 29; 44(2): e11

McCaffrey J, Young E, Lassahn K, Sibert J, Pastor S, Riethman H and Xiao M (2017) High-throughput single-molecule telomere characterization. Genome Res. 2017. 27: 1904–1915

Nowoshilow S, Schloissnig S et al. The axolotl genome and the evolution of key tissue formation regulators. Nature 554, 50–55

Qi LS, Larson MH, Gilbert LA, Doudna JA, Weissman JS, Arkin AP, Lim WA. (2013) Repurposing CRISPR as an RNA-guided platform for sequence-specific control of gene expression. Cell. 2013 Feb 28; 152(5):1173–83.

Sternberg SH, Doudna JA (2015) Expanding the Biologist’s Toolkit with CRISPR-Cas9. Mol Cell. 58(4):568–574

Tslil G, Sharim H et al. (2018) Epigenetic Optical Mapping of 5-Hydroxymethylcytosine in Nanochannel Arrays. ACS Nano doi: 10.1021/acsnano.8b03023

